# Identification of Memory Reactivation during Sleep by EEG Classification

**DOI:** 10.1101/200436

**Authors:** Suliman Belal, James Cousins, Wael El-Deredy, Laura Parkes, Jules Schneider, Hikaru Tsujimura, Alexia Zoumpoulaki, Marta Perapoch, Penelope Lewis

**Author notes:** **Corresponding Author:** Suliman Belal. School of Biological Sciences, Division of Neuroscience and Experimental Psychology, Manchester University, Zochonis Building, Brunswick Street, Manchester M13 9PT, UK.

## Abstract

Memory reactivation during sleep is critical for consolidation, but also extremely difficult to measure as it is subtle, distributed and temporally unpredictable. This article reports a novel method for detecting such reactivation in standard sleep recordings. During learning, participants produced a complex sequence of finger presses, with each finger cued by a distinct audio-visual stimulus. Auditory cues were then re-played during subsequent sleep to trigger neural reactivation through a method known as targeted memory reactivation (TMR). Next, we used electroencephalography data from the learning session to train a machine learning classifier, and then applied this classifier to sleep data to determine how successfully each tone had elicited memory reactivation. Above chance classification was significantly higher in slow wave sleep than in stage 2, suggesting differential efficacy of TMR in these two sleep stages. Interestingly, classification success reduced across numerous repetitions of the tone cue, suggesting either a gradually reducing responsiveness to such cues or a plasticity-related change in the neural signature as a result of cueing. We believe this method will be invaluable for future investigations of memory consolidation.

## Introduction

Newly learned memories are reactivated in sleep at both neuronal (Wilson and McNaughton, 1994; Jones and Wilson, 2005; Ego-Stengel and Wilson, 2007, 2010) and systems levels (Maquet et al., 2000; Peigneux et al., 2004). Such reactivation can be intentionally triggered through targeted memory reactivation (TMR), in which cues associated with previous learning are used to reactivate aspects of this prior learning on demand (Rasch et al., 2007; Rudoy et al., 2009; Diekelmann et al., 2011; Antony et al., 2012; Fuentemilla et al., 2013; Oudiette and Paller, 2013; Cousins et al., 2014, 2016; Schreiner and Rasch, 2014). Several influential models, including Active Systems Consolidation (Rasch and Born, 2013), Synaptic Homeostasis (Tononi and Cirelli, 2006, 2014), Memory Triage (Stickgold and Walker, 2013), and Information Overlap to Abstract (Lewis and Durrant, 2011), have proposed mechanisms by which memory reactivation in sleep could boost memory consolidation, but these ideas have been difficult to test since reactivation is notoriously problematic to detect in humans. The challenge stems both from not knowing precisely when during sleep reactivation occurs, and from the fact that reactivation can be greatly compressed in time (Nádasdy et al., 1999).

Prior attempts to measure reactivation in humans (Maquet et al., 2000; Peigneux et al., 2004) have provided evidence that neural activity during sleep partially mimics the activity occurring during wake, and that the extent of such reactivation can predict the degree of behavioural improvement across retention periods (Peigneux et al., 2004; Yotsumoto et al., 2009). Other work has used multivariate classification to capture the distributed signals associated with wakeful memory reactivation in functional magnetic resonance imaging (fMRI) (Deuker et al., 2013; Staresina et al., 2013) and magnetoencephalography (Fuentemilla et al., 2010). One fMRI study applied TMR in sleep to control the time at which reactivation occurred (van Dongen et al., 2012). More recent work has shown that electroencephalography (EEG) classifiers can distinguish between the sleep following two different learning tasks (Schönauer et al., 2017). In the current report, we build on these ideas by combining machine learning and TMR to control the timing of reactivation. Our goal is to identify neural reactivation on a trial by trial basis. We use EEG because of its excellent temporal resolution and appropriateness for sleep studies.

Our participants performed a sleep sensitive serial reaction time task (SRTT) (Cousins et al., 2014, 2016; Schönauer et al., 2014). Participants were intensively trained on a fixed sequence of finger presses cued by audio-visual triggers. To minimise motion artefacts, they were re-exposed to the audio-visual cues and asked to imagine making the cued movement while remaining motionless (‘Imagery task’). During subsequent sleep, we re-played cue tones to trigger reactivation of the associated memory. EEG data from the Imagery task were then used to train a multivariate classifier which was applied to the Sleep data to detect TMR cued reactivations.

Because much of the work on memory reactivation in sleep has been done in rats where stage 2 sleep (N2) and slow wave sleep (SWS) are not considered separately (Lee and Wilson, 2002; Ego-Stengel and Wilson, 2007; Bendor and Wilson, 2012), it remains unclear whether reactivation has distinct characteristics in these two sleep stages. Although TMR has been applied in both stages in humans (Rasch et al., 2007; Rudoy et al., 2009; Antony et al., 2012), no direct comparison has been made. We addressed this question by triggering reactivation in N2 and SWS, and examining the classification rate in both. It is currently unclear whether TMR always triggers reactivation, or whether the system eventually saturates, such that no further processing will occur. We tested whether the classification rate changed systematically across repeated TMR cues.

## Materials and Methods

The experiment was approved by the University of Manchester Ethics committee. Participants provided informed consent and were reimbursed for their time. 30 healthy volunteers with no self-reported history of neurological, psychiatric, sleep, or motor disorders participated, 15 (6 males, 27 ± 8 years) in the main experiment, and 15 (2 male, 25 ± 5 years) in the Control. All participants abstained from caffeine and alcohol for 24 hours prior to the experiment.

### Design and Procedure for the Main Overnight Experiment

Participants completed the Stanford Sleepiness Scale (Hoddes et al., 1972) at the start of each testing session (e.g. pre- and post-sleep). Participants were fitted for polysomnographic (PSG) recording at 8-9 pm before performing an adapted SRTT (Nissen and Bullemer, 1987) containing repeating blocks of a single fixed 12-item sequence (1-2-1-4-2-3-4-1-3-2-4-3). They were then permitted to read in bed until ~11 pm, and allowed to sleep for ~8 hours until 7-8 am (Figure 1A). During the night, tones associated with the learned sequence were softly played in blocks of repeating correct-order sequences during SWS and N2. Brown noise was played throughout the night to minimise disturbance.

**Figure 1.**
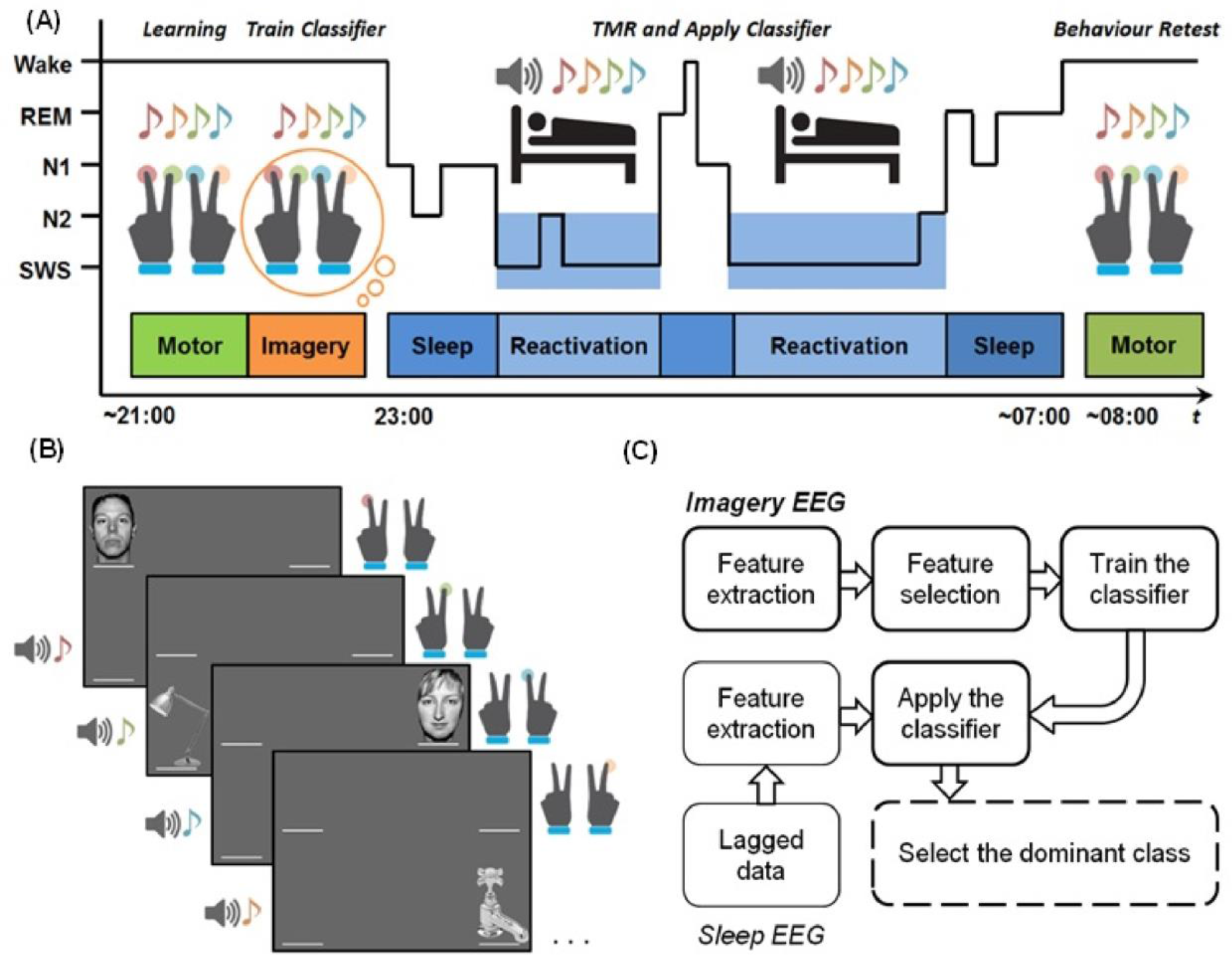
A schematic illustration of the design of the experiment and the classifier. **(A)** In the Motor task, participants performed the SRTT task with finger presses. In the Imagery task they were instructed to remain motionless and imagine performing the task while experiencing the same audio-visual cues that were used in the Motor task. During subsequent SWS and N2, the sequence was repeatedly reactivated in blocks of 1.5 min on, 2 min off. **(B)** The visual cues used in the experiment. Visual cues were objects or faces (1=face #1, 2=lamp, 3=face #2, 4=water tap). **(C)** The bold lines indicate processing the Imagery data, the thin lines are for the Sleep data and the dashed lines are for both Sleep and Imagery. Note that features were extracted from both Sleep EEG and Imagery EEG, but the classifier was trained on features from the Imagery data, and then applied to both the Sleep data and the Imagery data. Thus the classifier was not trained on the Sleep data.

For each trial in the SRTT, a visual cue appeared with a tone in one of four spatial locations, corresponding to keyboard keys of the same configuration, and participants pressed as quickly as possible while minimising errors with index and middle fingers. Items appearing on the left of the screen (Classes 1 and 2) required a left hand response, while items on the right side of the screen (Classes 3 and 4) required right hand responses. This arrangement was chosen to make the sequence easier to classify, as left handed responses are associated with a right hemispheric EEG response, and vice versa.

Visual cues were objects or faces (1=face #1, 2=lamp, 3=face #2, 4=water tap), see Figure 1B. Like the button response fingers, these images were chosen to make the sequence more easily classifiable since we expected the N170 component (Eimer, 2011) to be different for faces (Classes 1 and 3) and objects (Classes 2 and 4) at electrodes near face-selective brain regions, P7 and P8 (Calder et al., 2010). Each visual cue was accompanied by a specific auditory tone which was also associated with the cued finger. Tones (each lasting 300ms and played through headphones at an intensity which participants found comfortable) were musical notes grouped closely within the 4^th^ (low) (C/D/E/F) octave. Training comprised 7 blocks, with 10 sequence repetitions per block giving a total of 70 sequence repetitions, and 210 trials for each of the four finger classes. After each response there was a 1,230ms inter-trial interval before the next cue started.

After completion of training, participants performed a block of Imagery task SRTT in which the audio-visual cues were presented exactly as they had been during Motor training, but participants remained immobile, simply imagining they were pressing each button when cued. The Imagery session comprised 7 blocks of 10 sequence repetitions, and the EEG data from the Imagery session, which was free of motion artefacts, was used to train and test our classifier. Tone onsets were 1,500ms apart.

We recorded event-related potentials (ERPs) from 70 sequences in both Motor and Imagery tasks in 15 experimental participants. Due to noise on the trial-marker channel, two participants had lower trial numbers, thus ERPs from only 50 sequences were extracted from the Motor task in one, while only 60 sequences were extracted from the Imagery task in another.

During subsequent stable SWS and N2 sleep, tones were played softly (approximately 48dB) in blocks of 5 sequence repetitions. Tones were spaced 1,500ms apart. Each reactivation block took 1.5 minutes (5 × 12 × 1,500ms), and was followed by 2 minutes without reactivation. Reactivation was paused immediately upon signs of changes in sleep stage or arousal.

We performed TMR during SWS (79±38 sequences) in all 15 participants, and during N2 (83±45 sequences) in only 14 of these participants due to an experimenter error. The number of tone sequences presented in sleep varied between participants due to differences in the length of sleep stages. The precise numbers of sequences presented to each participant before sleep, during SWS or N2, and post-sleep is shown in **Table 1**.

**Table 1.** The number of 12-item sequences used for each participant before sleep, during sleep and after waking. Event-related potentials (ERPs) from 70 sequences were recorded in both Motor and Imagery tasks in 15 experimental participants. Due to noise on the trial-marker channel two participants had lower trial numbers, thus ERPs from only 50 sequences were extracted from the Motor task in one, while only 60 sequences were extracted from the Imagery task in another.

| Participant | Number of Presented Sequences |  |  |  |  |
| --- | --- | --- | --- | --- | --- |
|  | Before Sleep |  | During Sleep |  | Morning (Wake) |
|  | Motor (Learning) | Imagery | SWS | N2 | Motor (Retest) |
| 1 | 70 | 70 | 41 | No data | 70 |
| 2 | 70 | 70 | 99 | 59 | 70 |
| 3 | 70 | 60 | 24 | 39 | 70 |
| 4 | 70 | 70 | 99 | 199 | 70 |
| 5 | 70 | 70 | 54 | 114 | 70 |
| 6 | 70 | 70 | 139 | 84 | No data |
| 7 | 70 | 70 | 39 | 59 | 70 |
| 8 | 70 | 70 | 74 | 64 | 70 |
| 9 | 70 | 70 | 79 | 124 | No data |
| 10 | 50 | 70 | 74 | 49 | 70 |
| 11 | 70 | 70 | 99 | 119 | No data |
| 12 | 70 | 70 | 99 | 24 | No data |
| 13 | 70 | 70 | 157 | 60 | 70 |
| 14 | 70 | 70 | 39 | 110 | 70 |
| 15 | 70 | 70 | 62 | 91 | 70 |

The Motor task was repeated in the morning after the sleep experiment for 11 of the 15 participants in order to provide a measure of behavioural plasticity across the sleep epoch. Note that we did not collect these data in the first 4 participants.

### Control task

To demonstrate that it was not possible to classify the brain activity associated with simply hearing tones in the absence of any procedural learning, fifteen volunteers who were naïve to the experiment and task listened to the tones associated with the Imagery task with no visual input. The tone sequences were presented 140 times, which is equivalent to the total number of sequences presented during both Motor and Imagery tasks together, with a timing equivalent to that in the main experiment.

### EEG recording and analysis

Scalp electrodes were attached according to the 10-20 system at sixteen standard locations: F3, F4, C5, C3, Cz, C4, C6, CP5, CP3, CP4, CP6, P7, Pz, P8, O1, O2, and all referenced to the combined mean of left and right mastoid. Left and upper electrooculogram, left and right electrooculogram, and a forehead ground electrode were also attached. Impedance <5 kΩ was verified at each electrode, and the digital sampling rate was 200 Hz throughout the experiment. Data were scored by a trained sleep researcher according to the AASM Manual (American Academy of Sleep Medicine, Westchester, IL).

The experimental paradigm was programmed in MATLAB 6.5 (The MathWorks Inc., Natick, MA, 2000) and Cogent 2000 (Functional Imaging Laboratory, Institute for Cognitive Neuroscience, University College London). Sounds were presented via Sony noise cancelling headphones during the learning session and via PC speakers during sleep reactivation.

## Classifier Analysis

We aimed to create an EEG classifier which could identify the neural activity associated with each of the 5 possible classes (one for each finger, and one for baseline EEG – or a failure to reactivate), and then apply this to the EEG data collected after each TMR tone (Figure 1C). To create the classifiers, we extracted specific features from the EEG obtained for each trial in both Motor and Imagery tasks. We then performed feature selection to reduce the dimensions of the data and to maximise classification accuracy of the weak signals embedded in noisy EEG data. We adopted a hybrid feature selection algorithm consisting of two stages. First, a filter mechanism stage ranked features based on joint mutual information (Yang and Moody, 1999). Next, a wrapper mechanism searched for the best subset of these features, maximising classification accuracy. We trained Linear Discriminate Classifiers (Heijden et al., 2004) and tested them using the Imagery task EEG recordings. We then applied the trained classifiers to the EEG recorded after each TMR event in sleep to determine whether it was possible to detect memory reactivation by correctly determining which finger press had been cued. We compared the mean classification rates for TMR applied in N2 and SWS. Finally, we examined the effect of repeated TMR upon classification rate. Each of these steps is described in more detail in the following.

### The Classification of Motor and Imagery EEG

Data from the Motor and Imagery tasks were analysed separately, but using an identical method. We segmented the EEG data into epochs of 1,500ms with stimulus onset at 500ms. Each epoch was baseline corrected by subtracting the mean of 500ms of pre-stimulus EEG from the remaining 1,000ms. As visual inspection showed that the averaged ERPs at different electrodes occurred during the first 400ms post-stimulus, we used the 400ms directly after each TMR cue for the analysis of that trial. Importantly, in addition to the four classes relating to the four finger cues we formed a fifth (null) class representing ‘no TMR cue’, using randomly chosen 400ms EEG segments from the 2-minute inter-block intervals. The trials of the fifth class were baseline corrected, as in the other four classes, by subtracting the mean of the 500ms preceding EEG. Each trial was assigned a label *ω*_*i*_, where *i* ∊{1,2,‥,5}. Trials were randomly divided into two sets: training **ℜ**_*training*_ (60%) and evaluation **ℜ**_*evaluation*_ (40%), with the number of trials for each class kept equal across sets to maintain balance during training and evaluation. We extracted three families of features from the training and evaluation sets to obtain a comprehensive description of the EEG data: discrete wavelet transform (DWT) features, spectral features, and time domain features (the down-sampled average EEG), as explained below.

### Discrete Wavelet Transform features

The discrete wavelet transform has many advantages over other conventional spectral methods for processing EEG signals. It provides an optimal resolution in both time and frequency domains, and the condition of signal stationarity is not a requirement (Graps, 1995). This latter advantage is important since the EEG exhibits a non-stationary behaviour in a variety of contexts (Krystal et al., 1999). Therefore, wavelet analysis using a Daubechies-4 (DB4) wavelet (Daubechies, 1988) was used to decompose the EEG data from each electrode into five different levels of approximation (A1-A5) and detail coefficients (D1-D5). The frequencies corresponding to different levels of decomposition are presented in **Table 2**, which shows that the frequency range of the detail coefficients at level 5 (D5) is within the theta range (4-8 Hz), D4 is within the alpha range (8-13 Hz),D3 is within the beta range (13-30 Hz) and D2 is within the low gamma band (25-50 Hz). To maintain a good signal-to-noise ratio, the analysis was limited to the detail coefficients of frequencies up to 50 Hz. Therefore, the detail coefficients at levels 2 to 5 extracted from each EEG channel were concatenated to form the DWT features vector:

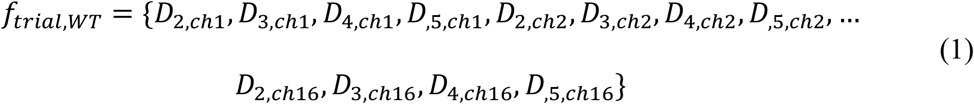

**Table 2.**
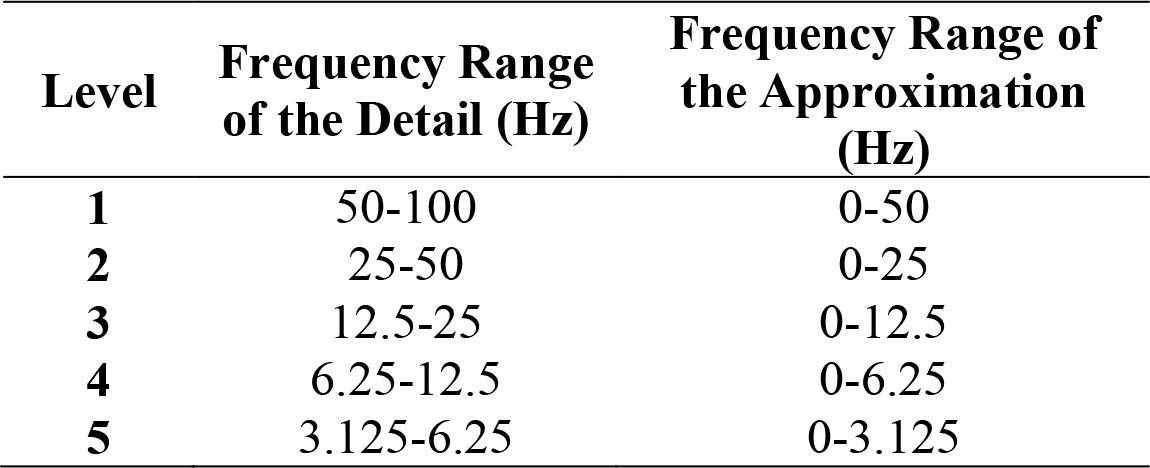
The frequencies corresponding to different levels of decomposition for Daubechies-4 (DB4) filter wavelet with a sampling frequency of 200 Hz.

| Level | Frequency Range of the Detail (Hz) | Frequency Range of the Approximation (Hz) |
| --- | --- | --- |
| 1 | 50-100 | 0-50 |
| 2 | 25-50 | 0-25 |
| 3 | 12.5-25 | 0-12.5 |
| 4 | 6.25-12.5 | 0-6.25 |
| 5 | 3.125-6.25 | 0-3.125 |

D1 was discarded as it represents a higher frequency band.

### Spectral features

The Power Spectral Density (PSD) techniques for spectral signal representation have been demonstrated to be robust and consistent for classification for Motor Imagery EEG data (Herman et al., 2008). We computed power spectra by using the Welch’s modified periodogram method (Welch, 1967), in which the EEG on each electrode was divided into overlapping segments each having 64 samples with an overlapping ratio of 90%. The segments were then weighted by a Hanning window function to reduce spectral leakage. Fourier transform was applied on the windowed segments to obtain the power density values, which were then averaged. The average power in the bands theta (4-8) Hz, alpha (8-12) and beta (16-24) Hz was obtained from a rectangle approximation of the integral of the signal’s PSD.

For each EEG trial, we concatenated the computed spectral average power values from each EEG channel to form the spectral features vector:

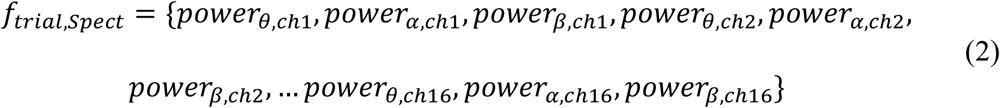

### Time-domain features

In order to include time domain information, we averaged the signal on each EEG channel (*EEG*_*ch*_) using a moving window of length *N* = 4.

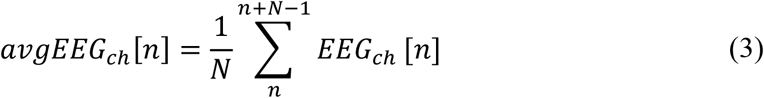

Next, we down-sampled the averaged EEG by a factor of four. For each EEG trial, we concatenated the down-sampled EEG to form the features vector:

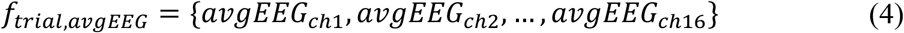

The EEG in **ℜ**_*training*_ and **ℜ**_*evaluation*_ was now represented by two feature matrices in which each row corresponds to a trial, and it consists of the concatenation of the DWT, spectral and time domain features. For each trial, 1,344 features were extracted (992 DWT features, 48 Spectral features and 304 Time-domain EEG features). Features in the training and evaluation sets were then normalised to zero mean and unit variance.

### Feature Selection

The values of each feature vector were transferred into three levels using quantile-based discretization. The features were then ranked using the joint mutual information method (Yang and Moody, 1999). The criterion for ranking features in this method provides the best trade-off in terms of accuracy, stability, and flexibility with small data samples (Brown et al., 2012).

Next, feature subsets were sequentially chosen in a forward manner from the ranked features, a normal-density-based linear discriminant classifier was trained using the features of each subset, and the classifier’s error rate was calculated. The initial subset contained the first two ranked features. New features were added to this initial subset if they led to a smaller error rate, but were otherwise discarded. This produced a monotonically decreasing error rate curve and an optimal feature subset. We used 10-fold cross validation (Kohavi, 1995) to evaluate each classifier. A normal-density-based linear discriminant classifier, which is widely used in EEG classifications, was used because it makes the posterior probabilities for each class available for further manipulation.

Once the optimal set of features for classification was chosen through feature selection, we calculated the performance of each classifier using **ℜ**_*evaluation*_. We used the correct classification rate (CCR) as a metric for evaluation. This was calculated as: (*N*_*Correct*_/ *N*_*Total*_) × 100 %. Where *N*_*Correct*_ is the number of correctly classified trials, and *N*_*Total*_ is the total number of trials to be classified.

In order to estimate the classification rate more robustly, we repeated the complete process five times, randomly selecting the training and evaluation sets each time. We then calculated the average evaluation classification performance and its standard deviation. Due to large inter-subject variability in the EEG, both feature selection and classifier training were conducted separately for each participant, such that each participant had their own individual classifier.

### Classification of sleep EEG

We first developed classifiers for the Motor task, where we expect movement related potentials to greatly facilitate classification. We tested these classifiers on 40% (held out) of the data. Next, we trained completely new classifiers on data from the Imagery task, as we expected these data to be more similar to what would be observed in sleep. We again tested these classifiers on 40% (held out) of the data. Next, we applied the Imagery Task classifiers to EEG data recorded during sleep to determine whether TMRs during sleep could be identified by classification. To do this, we performed an analysis similar to that used when applying classifiers to the held out Motor and Imagery data. However, in Sleep data, due to uncertainty about the timing of reactivation after the tone, the extraction process was repeated *n* = 120 times using a sliding window of length *W* = 400ms, and step size 5ms in order to maximise the chance of capturing a reactivation that could occur at any time during the 1,000ms after the tone. We then normalised the features extracted from the data. Finally, we applied the trained Imagery classifier to the normalised features extracted from each lag time, and obtained the class associated with each lag.

As a result, for each trial we obtained a vector containing *n* = 120 values for the class prediction (class label), each being: 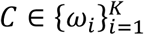, where *K* = 5. In a modified majority voting strategy, the class label with longest uninterrupted run, based on the process described below, was chosen as the predicted class of that trial.

As the response to the stimuli may occur at different points of time post-stimuli, we sought an optimal window for voting. We used 12 windows (subsets) of the 120 lags, shifting the start of each window by 10 lags. Thus, the first window contained the lags 1-120, the second window 10-120, the third 20-120, and the 12^th^ window contained the lags 110-120.

For each participant, this process was applied to the entire data set (all sequences). We then chose the window which corresponded to the highest classification rate across all trials for each participant as our classification window. Because classes were never repeated adjacently (e.g. class 1, class 1) in the trained sequence, classification predictions were constrained such that if such a repetition was chosen initially, the classifier was forced to choose the second-choice class, e.g. the class with the next longest run within the window.

To determine whether classification was above chance, we used a permutation test (Hesterberg et al., 2010; Lehmann and Romano, 2014). This consists of randomly shuffling the labels of the sleep trials and then calculating a new classification rate (‘random CCR’) for the shuffled data using the same set of features as selected before. The classification rate using the true labels, as calculated by sampling 50% of the data 1000 times and obtaining the mean, was compared against the ‘random CCR’ rates. A *p*-value for each participant was determined by counting the number of times that there was higher (or equal) classification accuracy in the shuffled data than in the true labels. Importantly, for each randomisation, we used the time window with the highest classification for the features of the shuffled labels, just as we had done for the true labels.

## Consistency of selected features

We sought to identify the most important features and electrodes by investigating the frequency of their selection (by the feature selection stage) at each of the 5 training sessions of the Imagery classifier.

To determine which were the most important down-sampled EEG features, the number of times each feature was selected across the 5 sessions for each participant was placed in a 2D-matrix, ‘feature selection matrix’, *X*(*n* × *m*), where *n* is the number of participants and *m* is the number of features. The columns of *X* were grouped separately into a multilevel cluster tree or “dendrogram” using hierarchical clustering. The Euclidian distance between pairs of features was calculated and the linkage criterion was the mean, in which the distance between two clusters is defined as the average distance between all pairs of the two clusters’ members. To determine which electrodes were most useful in classification, we applied the same approach by replacing the variables representing the rate of appearance of the features, *X*, by the frequency of the appearance of the electrodes.

We calculated the rates with which the four frequency bands of the DWT (25-50, 12.5-25, 6.25-12.5 and 3.125-6.25 Hz) were selected across participants, and then used a Friedman test to check for differences. Pairwise comparisons then established which bands were most useful for classification.

We followed a similar approach for spectral feature bands and for the type of features (feature families) selected by the training sessions. We also repeated the above analysis for the control data to see if there was any difference in the selected features.

## Control group

It was important to establish that the classifier was identifying the reactivation of memories associated with the stimuli and not just classifying auditory responses to the tones. In order to test this, we recorded data from 15 participants who were not aware of the underlying task or the purpose of the experiment. Participants listened to the same tones as had been presented in the ‘Motor’ and ‘Imagery’ tasks but without any visual stimuli or motor response. We then applied the classification analysis to the second block (equivalent to ‘Imagery’) in exactly the same way as it had been applied to the data collected from the original Imagery and Motor task. A total of 1680 trials were recorded from each of the 15 Control participants. From the second block of exposure, 60% of the trials were used for training and 40% for evaluation. We repeated the process of randomly sampling these percentages and training/testing the classifier and calculating the CCR 5 times (same as in the Experimental condition for the Imagery and Motor tasks) in order to provide a distribution of results for each participant.

To determine whether the CCR was above chance, we created a ‘random CCR’ by randomly shuffling the labels of the 40% of the Control data that was designated for test 1,000 times and then applying the 5 trained classifiers described above. We then compared the 5 random CCR results with the average value of the 5 CCR results for the correctly labelled trials to determine if classification in the latter was above chance, using the same method described above for classifying the sleep EEG.

## Behaviour

Our primary behavioural measure was the composite score (CS) of both response times (RT) and accuracy, CS = RT/accuracy (Bruyer and Brysbaert, 2011; Jackson et al., 2015), which was calculated using the mean values of each block of 10 sequences. Paired sample *t*-tests were used for the comparisons, except when the Shapiro-Wilk tests indicated a non-normal distribution, in which case Wilcoxon signed-rank tests were used. To minimise the contribution of outliers, we eliminated the trials in which the RT fell outside the mean± 2SD.

To determine whether there was a relationship between initial learning and the extent of reactivation in response to TMR (as determined by the CCR), we calculated a measure of initial learning strength by taking the difference between the first and last blocks of the Motor SRTT pre-sleep. We further investigated the relationship between the overnight improvement and the extent of reactivation. Overnight improvement was quantified using the difference between the last block of Motor SRTT pre-sleep and the first block of Motor SRTT post-sleep. We studied the above correlations during SWS and N2.

## Results

### Sleep measures

Polysomnography showed normal sleep architecture, with mean durations (minutes) in each sleep stage as follows: N1: 25.5 (21.6); N2: 253.4 (68.4); N3: 70.3 (32.0); REM: 70.9 (24.9); Wake: 35 (33.2), total sleep time 420 (35), and a mean sleep efficiency of 92.2 (8.2)%. Stanford sleepiness scores showed no difference in sleepiness levels between morning 3.11 (0.78) and evening 3.67 (1.66) sessions (*p* = 0.262, Wilcoxon signed-rank test). Note that polysomnographic data were lost for one participant, and Stanford Sleepiness Scale data were lost for 3.

### Classification of Motor and Imagery EEG

Both Motor and Imagery classifiers categorised trials at a high correct classification rate (CCR) of 0.70± 0.12 (mean ± SD) for Motor and 0.57 ± 0.16 for Imagery, (Figure 2A). Note that chance was 0.20 due to the five possible classes. CCRs for each participant are shown in Table 3. To ensure that this mean classification rate was not driven by detection of the fifth (‘no cue’) class, we separately examined mean classification of the four tone-related classes, which was well above chance with a CCR of 0.67 ± 0.15 for Motor and 0.54 ± 0.18 for Imagery. Unsurprisingly, classification of the fifth class was even higher, with a CCR of 0.82 ± 0.10 for Motor and 0.67 ± 0.15 for Imagery, indicating that our classifier can very successfully determine whether or not a tone was present.

**Figure 2.**
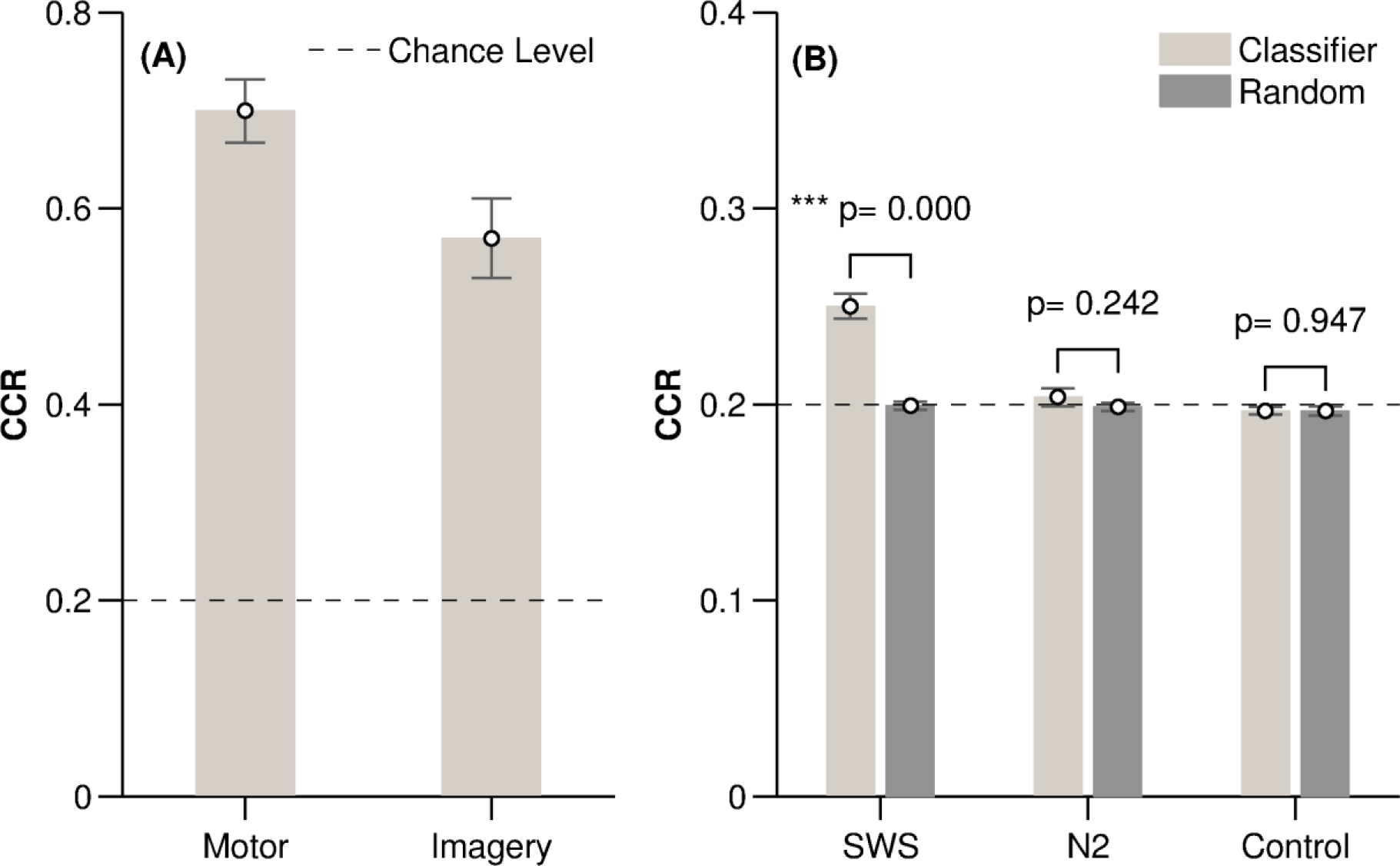
Behavioural results. (A) Correct classification rate (CCR) in the Motor and Imagery experiments shown as mean and standard error (SE). (B) Correct classification rate for SWS, N2 and Control and their corresponding random classifiers, shown as mean and SE.

**Table 3.**
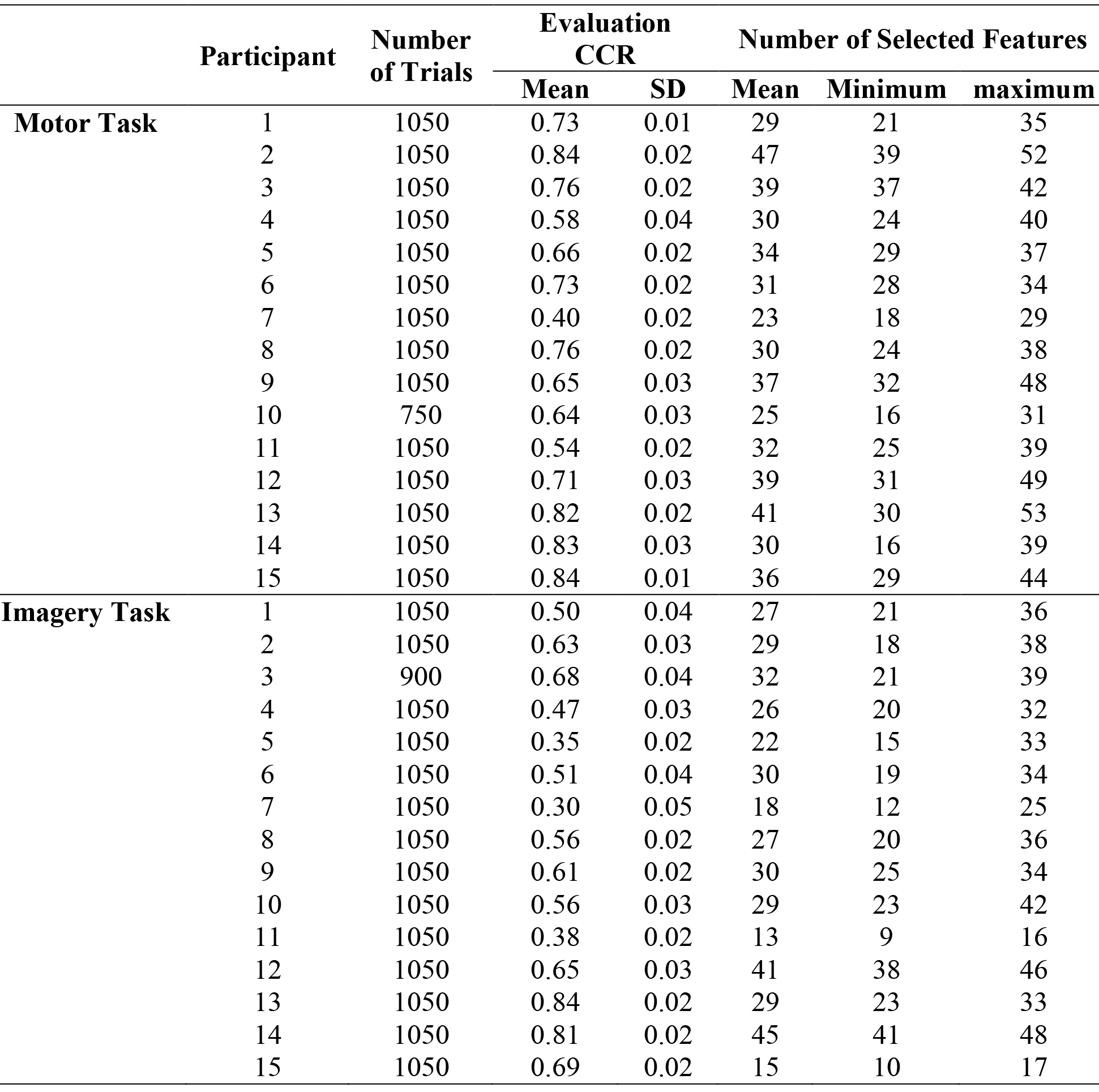
Motor and Imagery tasks classification. The average correct classification rate (CCR) of the evaluation ERP data for the Motor and Imagery tasks. Classifiers were trained using randomly selected subsets (60% of the data) and this was repeated 5 times. The trained classifiers were applied on the unseen evaluation data (40% of the data). SD: is the standard deviation over the 5 repeats.

|  | Participant | Number of Trials | Evaluation CCR |  | Number of Selected Features |  |  |
| --- | --- | --- | --- | --- | --- | --- | --- |
|  |  |  | Mean | SD | Mean | Minimum | maximum |
| **Motor Task** | 1 | 1050 | 0.73 | 0.01 | 29 | 21 | 35 |
|  | 2 | 1050 | 0.84 | 0.02 | 47 | 39 | 52 |
|  | 3 | 1050 | 0.76 | 0.02 | 39 | 37 | 42 |
|  | 4 | 1050 | 0.58 | 0.04 | 30 | 24 | 40 |
|  | 5 | 1050 | 0.66 | 0.02 | 34 | 29 | 37 |
|  | 6 | 1050 | 0.73 | 0.02 | 31 | 28 | 34 |
|  | 7 | 1050 | 0.40 | 0.02 | 23 | 18 | 29 |
|  | 8 | 1050 | 0.76 | 0.02 | 30 | 24 | 38 |
|  | 9 | 1050 | 0.65 | 0.03 | 37 | 32 | 48 |
|  | 10 | 750 | 0.64 | 0.03 | 25 | 16 | 31 |
|  | 11 | 1050 | 0.54 | 0.02 | 32 | 25 | 39 |
|  | 12 | 1050 | 0.71 | 0.03 | 39 | 31 | 49 |
|  | 13 | 1050 | 0.82 | 0.02 | 41 | 30 | 53 |
|  | 14 | 1050 | 0.83 | 0.03 | 30 | 16 | 39 |
|  | 15 | 1050 | 0.84 | 0.01 | 36 | 29 | 44 |
| **Imagery Task** | 1 | 1050 | 0.50 | 0.04 | 27 | 21 | 36 |
|  | 2 | 1050 | 0.63 | 0.03 | 29 | 18 | 38 |
|  | 3 | 900 | 0.68 | 0.04 | 32 | 21 | 39 |
|  | 4 | 1050 | 0.47 | 0.03 | 26 | 20 | 32 |
|  | 5 | 1050 | 0.35 | 0.02 | 22 | 15 | 33 |
|  | 6 | 1050 | 0.51 | 0.04 | 30 | 19 | 34 |
|  | 7 | 1050 | 0.30 | 0.05 | 18 | 12 | 25 |
|  | 8 | 1050 | 0.56 | 0.02 | 27 | 20 | 36 |
|  | 9 | 1050 | 0.61 | 0.02 | 30 | 25 | 34 |
|  | 10 | 1050 | 0.56 | 0.03 | 29 | 23 | 42 |
|  | 11 | 1050 | 0.38 | 0.02 | 13 | 9 | 16 |
|  | 12 | 1050 | 0.65 | 0.03 | 41 | 38 | 46 |
|  | 13 | 1050 | 0.84 | 0.02 | 29 | 23 | 33 |
|  | 14 | 1050 | 0.81 | 0.02 | 45 | 41 | 48 |
|  | 15 | 1050 | 0.69 | 0.02 | 15 | 10 | 17 |

### Classification of Sleep EEG

We applied the classifier that had been trained on Imagery task data to one second of EEG after each TMR tone in sleep. Due to uncertainty about when reactivation occurs after the TMR cue, we repeated feature extraction 120 times using a sliding window of 400ms. In a modified majority voting strategy, the class label with the longest uninterrupted run over the 120 extractions was chosen as the predicted class of that trial.

In SWS, classification was significantly above chance in all 15 participants (*t*(14) = 7.91, *P* < 0.0005), with a group mean CCR of 0.25 ± 0.03. In N2, classification was more variable, with above chance performance in only 5 of the 14 participants with TMR applied in N2, and a group mean of 0.20 ± 0.02 which did not differ from chance (*t*(13) = 0.79, *p* = 0.44); see **Table 4**.

**Table 4.** Sleep (SWS and N2) classification rates. Statistical comparisons between the mean correct classification rate (CCR) of the the TMR cued reactivations during sleep (SWS and N2) and the CCR of a random classifier. The mean and standard deviation of the classifier’s CCR were calculated after sampling the data (50%) 1000 times. For the random classifier, the class-labels were randomly shuffled before sampling. The start of the window corresponds to the sample index at which the optimal window for voting was chosen (see the materials and methods section). Cases in which no above chance classifier was found are indicated by ‘†’.

| Sleep Stage | Participant | The start of the window (ms) | CCR ± SD | | *p*-value |
| --- | --- | --- | --- | --- | --- |
|  |  |  | Classifier | Random Classifier |  |
| SWS | 1 | 50 | $0.28 \pm 0.02$ | $0.21 \pm 0.02$ | 0.004 |
| | 2 | 550 | $0.24 \pm 0.01$ | $0.20 \pm 0.01$ | 0.002 |
| | 3 | 400 | $0.25 \pm 0.02$ | $0.19 \pm 0.02$ | 0.017 |
| | 4 | 1 | $0.27 \pm 0.01$ | $0.20 \pm 0.01$ | $< 0.001$ |
| | 5 | 1 | $0.24 \pm 0.02$ | $0.19 \pm 0.02$ | 0.018 |
| | 6 | 1 | $0.28 \pm 0.01$ | $0.21 \pm 0.02$ | $< 0.001$ |
| | 7 | 450 | $0.21 \pm 0.02$ | $0.19 \pm 0.02$ | 0.040 |
| | 8 | 450 | $0.24 \pm 0.02$ | $0.19 \pm 0.02$ | 0.004 |
| | 9 | 200 | $0.26 \pm 0.01$ | $0.21 \pm 0.01$ | $< 0.001$ |
| | 10 | 300 | $0.25 \pm 0.01$ | $0.21 \pm 0.02$ | 0.016 |
| | 11 | 550 | $0.22 \pm 0.01$ | $0.20 \pm 0.01$ | 0.030 |
| | 12 | 550 | $0.23 \pm 0.01$ | $0.19 \pm 0.02$ | $< 0.001$ |
| | 13 | 500 | $0.23 \pm 0.01$ | $0.20 \pm 0.1$ | 0.02 |
| | 14 | 100 | $0.30 \pm 0.02$ | $0.20 \pm 0.03$ | 0.001 |
| | 15 | 100 | $0.25 \pm 0.02$ | $0.20 \pm 0.03$ | 0.011 |
| N2 | 1 | *No data* |  |  |  |
| | 2 | 500 | $0.21 \pm 0.01$ | $0.20 \pm 0.02$ | 0.070† |
| | 3 | 500 | $0.22 \pm 0.02$ | $0.21 \pm 0.01$ | 0.043 |
| | 4 | 500 | $0.22 \pm 0.01$ | $0.20 \pm 0.02$ | 0.004 |
| | 5 | 50 | $0.21 \pm 0.01$ | $0.19 \pm 0.01$ | $< 0.001$ |
| | 6 | 80 | $0.19 \pm 0.01$ | $0.20 \pm 0.02$ | 0.060† |
| | 7 | 100 | $0.18 \pm 0.01$ | $0.21 \pm 0.01$ | $< 0.001$† |
| | 8 | 50 | $0.21 \pm 0.02$ | $0.20 \pm 0.01$ | 0.039 |
| | 9 | 110 | $0.20 \pm 0.01$ | $0.20 \pm 0.02$ | 0.131† |
| | 10 | 110 | $0.16 \pm 0.02$ | $0.18 \pm 0.03$ | $< 0.001$† |
| | 11 | 80 | $0.20 \pm 0.01$ | $0.20 \pm 0.02$ | 0.245† |
| | 12 | 550 | $0.21 \pm 0.02$ | $0.19 \pm 0.02$ | 0.044 |
| | 13 | 450 | $0.21 \pm 0.02$ | $0.20 \pm 0.02$ | 0.316† |
| | 14 | 450 | $0.22 \pm 0.01$ | $0.20 \pm 0.02$ | 0.085† |
| | 15 | 550 | $0.21 \pm 0.01$ | $0.20 \pm 0.02$ | 0.241† |

To ensure that above-chance classification rates were not driven by class 5, we examined classification of the four tone-related classes, which were again well above chance in SWS, with a CCR 0.23 ± 0.02 > 0.20, *t*(14) = 6.64, *p* < 0.0005). CCR for the fifth class was also high (0.28 ± 0.13 > 0.20, *t*(14) = 2.38, *p* = 0.032 < 0.05). In N2, the figure was 0.19 ± 0.03 < 0.20, *t*(13) = −7.74, *p* = 0.427 for the four tone classes combined, and (0.27 ± 0.09 > 0.20, *t*(13) = 2.684, *p* = 0.019 < 0.05) for the fifth class.

As a further control, we compared the CCR from SWS and N2 with a ‘random CCR’, created by shuffling the trial labels. This showed greater group mean CCR for SWS (*t*(14) = 10.79, *p* < 0.0005), but not for the N2 classifiers (*p* = 0.242, Wilcoxon signed-rank test) when compared to random, Figure 2B and **Table 4**. However, the group mean CCR of above-chance N2 classifiers was different from random CCR (*p* = 0.038, Wilcoxon signed-rank test).

Furthermore, CCRs showed greater classification success in SWS than N2, both when all N2 participants were included (*t*(13) = 6.464, *p* < 0.0005), and when only above-chance classifier N2 participants were considered (*p* = 0.039, Wilcoxon signed-rank test).

### Classification of Control stimuli

Fifteen Control participants listened to the same auditory sequence as experimental participants, but without having learned any association between these and visual display or movement. Classification of the four tones was at chance level (CCR = 0.20 ± 0.01, *t*(14) = −1.31, *p* = 0.211), indicating that our classifier cannot discriminate between these tones unless associated with other information. As expected, CCR of the fifth class was above chance, (0.55 ± 0.01 > 0.20, *t*(14) = 14.02, *p* < 0.0005), indicating that the classifier can successfully detect the presence of a tone.

We also performed a second control analysis by comparing the CCR rates for the 4 tones and the background EEG with no tone presented to CCRs generated by randomly shuffling the trial labels. Permutation tests showed no difference between the correctly labelled and the random classifier in any Control participant, (*p* > 0.05 in all 15 cases). This result was also supported by a group comparison of the CCR for the four tones and their corresponding random CCRs confirmed (*t*(14) = 0.068, *p* = 0.947, Figure 2B). This again demonstrates that our classifier could not discriminate between the four tones unless they were associated with learned material.

Notably, the random CCR of the fifth class (background EEG) was artificially inflated through a bias towards the classification of all trials as background, and was thus above chance (0.28 ± 0.02 > 0.20, *t*(14) = 14.415, *p* < 0.0005). Irrespective of this artificial boosting, the random CCR was still significantly lower than the CCR of the control EEG, so despite this response bias, the classifier could discriminate between EEG and tone presentation.

### Consistency of features and electrodes used for classification

To determine which features were most useful for classification, we asked how often each feature in each of the three families of features (DWT features, spectral features, and time domain features) was selected by the feature selection stage. Feature selection rates were then compared both between and within families. This was repeated for the Imagery classifier and Control classifiers.

The selection rates of the three families of features differed significantly: Imagery classifier Friedman’s χ^2^(2, N = 15) = 24.13, *p* < 0.001 and Control-trained classifier, Friedman’s χ^2^(2, N = 15) = 26.27, *p* = 0.001 < 0.05, and all possible pairs of families differed from each other: Imagery post-hoc Wilcoxon *p* < 0.05, and Control-trained classifier post-hoc Wilcoxon *p*<0.05.

Interestingly, in the Imagery classifier, which easily distinguished between the four finger classes, the DWT features were consistently the most commonly selected. In the Control classifier, which could only distinguish between presence and absence of a tone, the down-sampled average EEG features were most commonly selected, see Figure 3. Furthermore, within the DWT family, there was no statistical difference in the selection of the coefficients of the different frequency bands (Friedman’s χ^2^(3, N = 15) = 6.3, *p* = 0.098). This was not the case for the coefficients selected by the Control classifier (Friedman’s χ^2^(3, N = 15) = 18.84, *p* < 0.001). The coefficients of higher frequencies (25-50 Hz) were more commonly selected in the Control classifier (Wilcoxon, *p* < 0.05) while the lower frequencies (3.125-6.25 Hz) were the least selected (Wilcoxon, *p* < 0.05).

**Figure 3.**
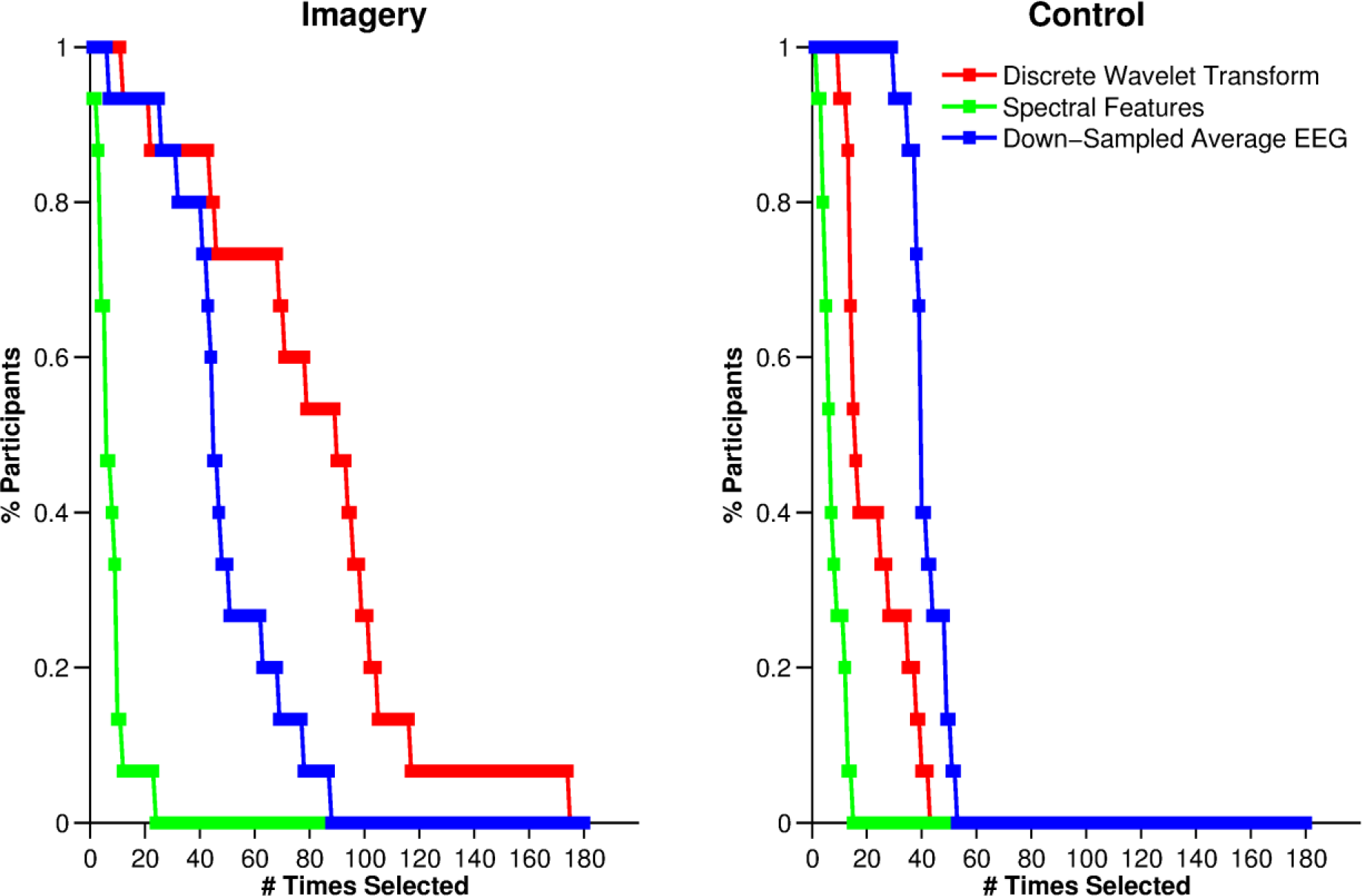
Frequency of selecting each family of features. After the feature extraction stage, a feature selection process determines which features were most suitable for classification. The X-axis (# Times Selected) represents the number of times each feature family appeared across participants. Y-axis (% participants) shows the proportion of participants in whom that particular number of features was selected.

Within the time domain family, the 19 features were selected at different rates in both Imagery and Control classifiers, Friedman’s χ^2^(18, N = 15) = 51.42, *p* < 0.001 and χ^2^(18, N = 15) = 82.95, *p* < 0.001, respectively. Hierarchical clustering showed that features 4, 5, 6, 7, 8, 10 and 12, were the most selected for the imagery trials while features 1 to 8 were the most selected for the control trials, with feature 8 being the most frequent in both. The time domain features 4, 5, 6, 7, 8, 9 and 10 represented the amplitudes of the elicited ERP in the interval 40-200ms which captured the P1 and N1 components of the ERP. Because the time domain feature family was consistently the most useful in the control classifier, and because that the classifier could detect the presence or absence of a tone but nothing more, this finding suggests that the ERP peaks were useful for such determinations.

Within the spectral features family, which was the least consistently used by both classifiers, both 4-8 and 8-12 Hz bands were most frequently selected in the Imagery classifier Friedman χ^2^(2, N = 15) = 5.35, *p* = 0.067 > 0.05 whereas only 4-8 Hz was frequently selected in the Control classifier (Friedman χ^2^(2, N = 15) = 27.1, p = 0.001 < 0.05; Wilcoxon, *p* < 0.0005).

To determine which electrodes out of our array of 16 provided the most useful information for classification, we repeated the above analysis now considering the electrodes at which features were selected. This revealed a consistent difference in the number of times specific electrodes were selected in both Imagery, Friedman’s χ^2^(15, N = 15) = 39.3, *p* < 0.001 and Control data, Friedman’s χ^2^(15, N = 15) = 58.29, *p* = 0.001 < 0.05. Hierarchical clustering showed that electrodes F3, F4, P7, P8,Pz and C5 (Figure 4), and F3, F4, C3, C4, C5, C6, Cz and P8 were the most frequently selected for the Imagery and Control trials, respectively. P8 was the most frequently selected in the Imagery trials and Cz in the Control trials.

**Figure 4.**
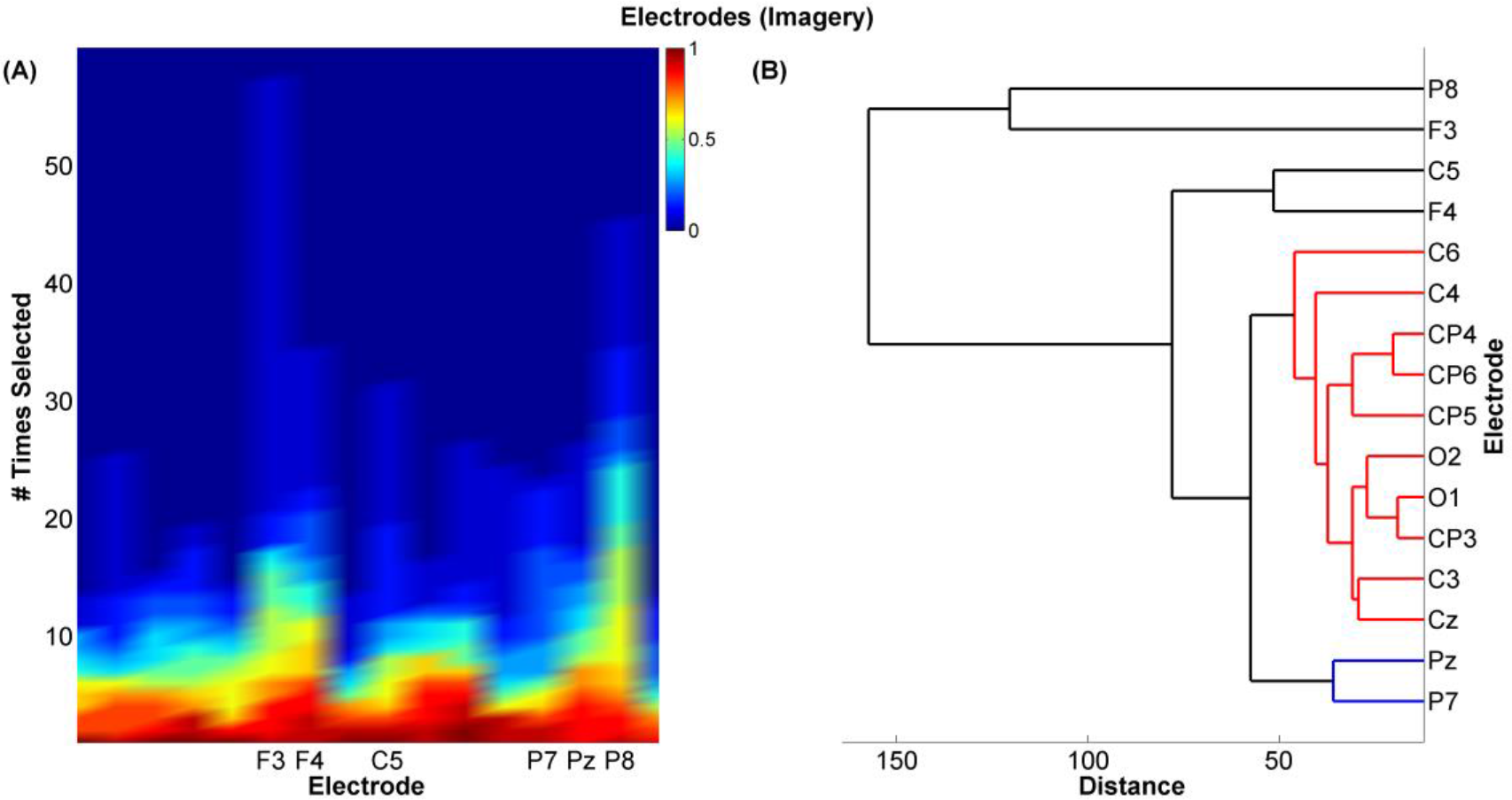
Electrode selection. **(A)** A plot of the frequency of selecting each of the 16 electrodes for the Imagery classifier. This was determined by accounting for each time a feature belonging to a particular electrode was selected by the classifier. The more often an electrode was selected (# Times Selected) across a large proportion of the participants (% Participants indicated by the colour bar), the more important the electrode was deemed. This was objectively determined using hierarchical clustering **(B)**.

### The effect of multiple TMR repetitions on classifier performance

We next set out to determine whether classification strength changed across repeated TMR events. We calculated classifier performance for each participant using a sliding window of 240 trials in length. This revealed a significant (*p* < 0.001) decrease in CCR across repetitions during SWS in 13 of 15 participants. In N2, decrease across TMR repetitions was significant (*p* < 0.001) in only 2 of the 5 participants who showed above chance classification, **Table 5**.

**Table 5.**
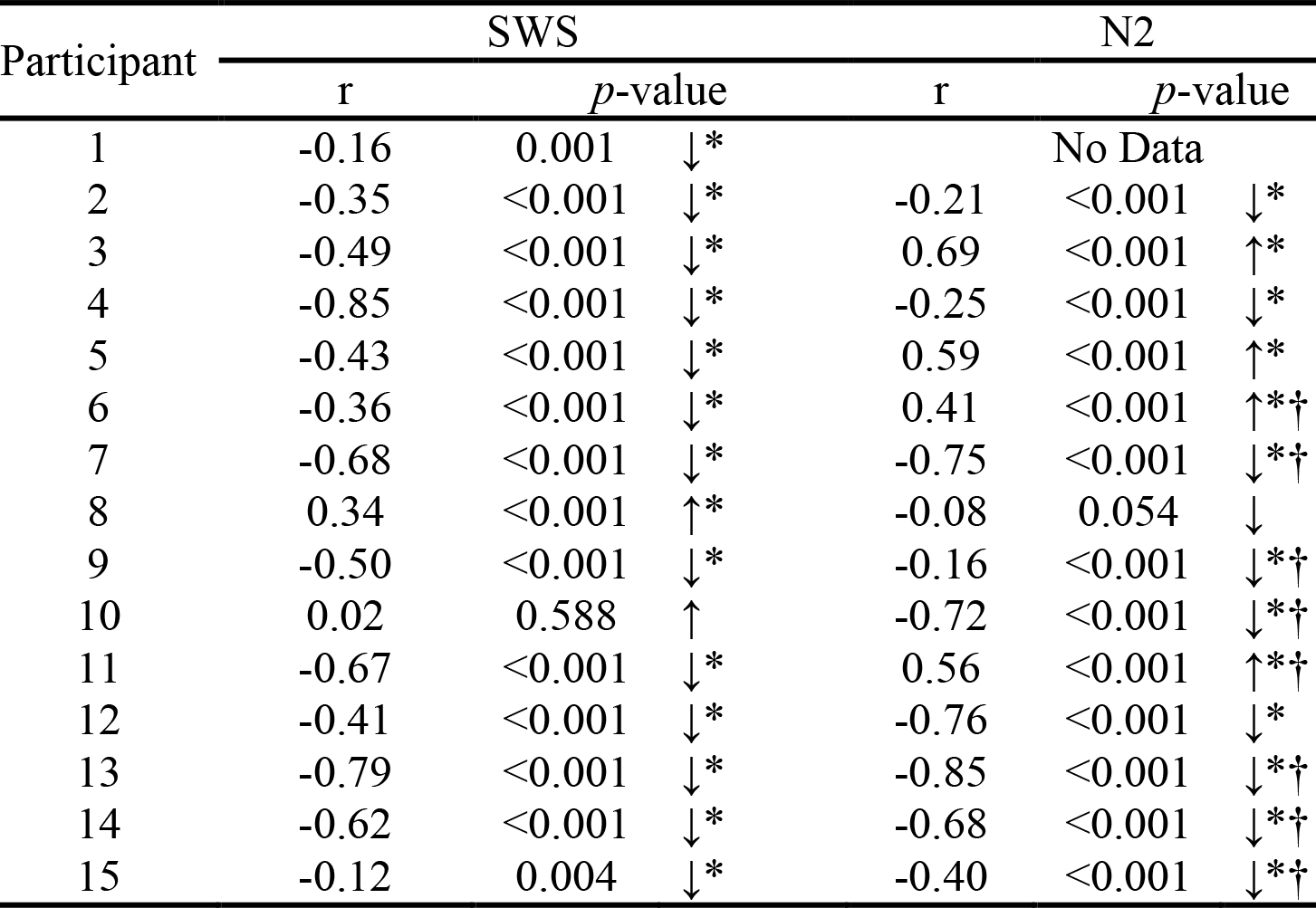
The correlation between the CCR and the repetitions during SWS and N2. The arrows indicate the direction (positive or negative) of the correlations, ‘*’ significant correlations (p < 0.05), and ‘†’ classifier is not above chance.

### Behaviour

We were interested in relationships between behavioural performance and CCR, and tested for these in the participants who completed the full behavioural task (before and after sleep). One participant was excluded due to performance decrease across training which indicated disengagement from the task. To determine whether stronger initial learning was associated with a greater classification rate for TMR cued replays during subsequent sleep, we computed a measure of initial learning by calculating the change in performance using composite score (CS = speed/ accuracy) between the first and last test blocks of the pre-sleep Motor task (Bruyer and Brysbaert, 2011; Jackson et al., 2015). A Pearson correlation between this measure and CCR during SWS revealed no significant relationship (r = −0.11, *p* = 0.769).

Examination of how performance changed across sleep revealed the expected improvement in CS (*t*(9) = 5.632, *p* < 0.001). However, Pearson correlations between CCR in SWS and this improvement revealed no significant relationship between CCR and the overnight change in CS (Pearson r = −0.42, *p* = 0.224).

Repeating the above behavioural correlation analyses in participants who completed the post-sleep behavioural tasks and also exhibited an above chance CCR in N2 showed no significant correlations between either initial learning or overnight improvement and N2 CCR. This could be due to the small sample size (n=5).

## Discussion

We have developed a non-invasive method for identification of neural reactivation in sleep, demonstrating as a proof of principle that it is possible to detect TMR cued reactivations above chance level in SWS using EEG classifiers. Through applying this method, we provide critical support for the occurrence of memory reactivation during human sleep, and for the triggering of such reactivation with TMR. We also show that repeated triggering of reactivation in SWS results in a gradually decreasing classification rate. Additionally, our behavioural data suggest that it is easier to detect the replay of memories that are better learned before sleep. Because our method uses EEG data, which is standardly recorded during sleep experiments, we believe it will provide a highly convenient tool for future examinations of memory consolidation in sleep.

### Classifiers

Our classification pipeline was specifically tailored to identification of TMR trials during sleep. We used an array of 16 electrodes; however, post-hoc analyses revealed that only a subset of these were consistently useful for classification. Because our task requires integration of visual, auditory, and motor information it seems plausible that the utility of parietal electrode P8 for classification of imagery trials in the majority of participants may be due to the cross-modal integration function of this area. We selected which families of features to include based on the nature of the EEG signals and the characteristics of the classes we were aiming to predict. EEG signals are non-stationary, and we had to consider the possibility that responses to TMR during sleep could be a compressed version of responses during wake. The coefficients of the wavelet transform, the spectral power, and the ERPs were therefore all potential candidate features; however, it was interesting to note that the wavelet transform and ERP families consistently provided useful information, while the spectral power did not. It is similarly noteworthy that the lower frequency information which characterises sleep was consistently useful in classification, while high frequency information was not.

### SWS vs N2

We observed a significantly higher classification rate in SWS than N2 although 5 out of 14 participants tested did exhibit classification significantly above chance in N2. The higher classification rate in SWS could suggest that although reactivation in N2 clearly occurs and can be triggered with TMR, it is qualitatively different from reactivation in SWS, and thus not as readily identifiable using the same classifier. The fact that EEG activity in SWS differs more markedly from wake (where the classifier was trained) than EEG in N2 means this is unlikely to be a signal to noise issue. Furthermore, the frequent slow oscillation troughs in SWS inhibit cortical activity, and can thus be expected to inhibit neural reactivation as well (Cox et al., 2014), reducing the success of TMR tones which were played without attention to these cycles. Because these troughs do not exist to the same extent in N2, if reactivation was equally strong in both stages we might expect the classification rate to be stronger in N2. The fact that TMR in N2 resulted in less robust classification than in SWS therefore suggests that TMR in N2 does not elicit reactivation to the same extent as TMR in SWS, a difference which might relate to the differential physiology of these two stages, e.g. different levels of acetylcholine and differential connectivity between the hippocampus and neocortex (Andrade et al., 2011).

### Decay of classification rate

Our observation that the rate of classification decays across repeated TMR applications in SWS can be interpreted in two different ways. First, once they have been reactivated a certain number of times in a night, memories may no longer be as likely to reactivate in response to TMR. This idea is in keeping with the observation that neural reactivation in rats declines sharply across the first hour of sleep (Tatsuno et al., 2006), and could occur because memories have been processed to a sufficient degree, see (Vyazovskiy and Delogu, 2014), or even to the maximal degree possible in one night. Alternately, our finding could suggest that the neural signature of reactivation evolves across TMR events, such that it eventually does not fit the classifier we developed before sleep, an explanation which could also be relevant for this effect in template-matching based reactivation studies in rats (Tatsuno et al., 2006). This latter idea builds on neuroimaging data (Walker et al., 2005; Takashima et al., 2006; Gais et al., 2007; Sterpenich et al., 2007; Durrant et al., 2012) showing that the neural signature of remembering is different after sleep, and this plasticity often relates to the amount of SWS obtained.

## In sum

We have developed a method for detecting neural reactivation in sleep using EEG classifiers. This will provide a useful tool for future explorations of such reactivation and its impacts on memory consolidation and brain plasticity. In the current proof of principle paper, we have applied this method to two specific problems. We show that while TMR elicits classifiable reactivations in both SWS than N2, these are more consistently classifiable in SWS. We also show that TMR induced reactivation becomes less classifiable with multiple repetitions suggesting that TMR becomes less effective as the neural processing associated with reactivation is gradually completed. In future, our classifier method could be applied to determine whether more classifiable reactivations lead to greater functional plasticity, and which EEG features are the most important for this.

## Acknowledgements

This work was supported by the Wellcome Trust Institutional Strategic Support Fund (grant number R117875) and by the University of Manchester.

